# Phosphoregulation of RAD51AP1 function in homology-directed repair

**DOI:** 10.1101/2025.09.10.675389

**Authors:** Neelam Sharma, Mollie Uhrig, Youngho Kwon, Patrick Sung, Claudia Wiese

## Abstract

Homology-directed DNA repair (HDR) is critical for genome stability and tumor suppression. HDR is initiated by the RAD51 single-stranded (ss)DNA nucleoprotein filament which conducts the homology search and invades a homologous DNA template, creating a displacement-loop (D-loop). The RAD51 filament is assisted in these processes by several proteins. One such protein is RAD51-Associated-Protein 1 (RAD51AP1) which binds DNA and RNA and directly interacts with RAD51. Of note, RAD51AP1 overexpression is associated with poor prognosis in several different cancer types.

Here, we show that RAD51AP1 activity is regulated by phosphorylation. RAD51AP1 bearing S277/282A mutations is more proficient in the stimulation of D-loop formation than wild type RAD51AP1 or phosphomimetic RAD51AP1-S277/282D. In EMSAs, RAD51AP1 with S277/282A mutations more avidly binds ssDNA, double-stranded (ds)DNA, and the nucleosome core particle than wild type RAD51AP1 or RAD51AP1-S277/282D. In cells, RAD51AP1-S277/282A confers no rescue of RAD51AP1 deficiency in toxicity tests and DNA replication assays. In contrast, RAD51AP1-S277/282D fully rescues RAD51AP1 deficiency. We provide evidence that RAD51AP1-S277 is a CDK2 target and propose a model in which RAD51AP1-S277/282 phosphorylation ensures RAD51AP1 flexibility for dynamic engagement in consecutive steps of the HDR reaction. Our results provide new mechanistic insights into RAD51AP1 regulation by a CDK.

## INTRODUCTION

Homology-directed DNA damage repair (HDR) is a critical tumor suppressor pathway that is defective in patients with BRCA1/2 mutations. HDR requires a homologous double-stranded (ds)DNA donor molecule to direct the repair of double-strand breaks (DSBs), inter-strand DNA crosslinks (ICLs), and other DNA lesions. HDR is initiated by 5’-3’end resection of the DSB to generate a 3’-single-stranded (ss)DNA overhang (1, 2). This 3’-ssDNA overhang is protected by the ssDNA-binding protein RPA. RPA is replaced by RAD51, a process that requires the action of multiple mediator proteins (3–7). The RAD51-ssDNA nucleoprotein filament then captures the homologous dsDNA donor and generates a displacement loop (D-loop) by strand invasion. In this process, the RAD51 filament is supported by several accessory HDR factors (8–13).

RAD51-Associated Protein 1 (RAD51AP1) is a critical RAD51-interacting protein that assists RAD51 during the strand invasion step in HDR (8, 9, 14). RAD51AP1 was identified as a RAD51-interacting protein in yeast two-hybrid library screens (15, 16), and its RAD51 interaction domain has been mapped to the 25 C-terminal residues in RAD51AP1 (8, 9, 17). RAD51AP1 contains two nucleic acid binding motifs to recognize nucleic acids, one of which is in its N-terminal domain and the second is in its C-terminal region (9, 18). As such, RAD51AP1 has high affinity for ssDNA, dsDNA, the D-loop structure, and the nucleosome core particle (NCP), and binds RNA avidly (8, 9, 15, 19, 20).

RAD51AP1 functions in the protection of cells from genotoxic agents (8, 9, 21). RAD51AP1 also functions in the HDR-mediated alternative lengthening of telomeres (ALT) pathway through its direct interaction with telomere repeat-containing lincRNA (22, 23). Mechanistically, RAD51AP1 promotes the invasion of RNA into DNA, promoting DR-loop formation (24). In a clinical context, RAD51AP1 expression is significantly upregulated in several different tumor types, and this increase is associated with resistance to therapy and poor prognosis (14, 25, 26). Yet, how the activity of the RAD51AP1 protein is regulated is poorly understood.

HDR relies on the undamaged sister chromatid to provide the template for repair. Thus, HDR is tightly coordinated with cell cycle progression (27, 28). Such coordination can be achieved by cyclin-dependent kinases (CDKs), which have been shown to target several critical proteins in the HDR pathway. For example, CDK-dependent phosphorylation of T847 in CtIP is required for the initiation of 5’-3’ resection (29, 30). The interaction between BRCA1 and CtIP depends on CtIP-S327 phosphorylation mediated by a CDK (31–33). CDK2 phosphorylates S276/T315 in CtIP to enable CtIP’s association with the cis-trans prolyl isomerase PIN1, leading to a downregulation of resection (34). CDK2 was also shown to phosphorylate UHRF1-S674, an E3 ubiquitin ligase, allowing BRCA1-mediated removal of RIF1 from sites of DNA damage (34, 35). The CDK-dependent phosphorylation of BRCA2-S3291, which regulates BRCA2’s C-terminal interaction with RAD51, has also been well characterized (36, 37).

Phospho-proteomic screens have identified phospho-peptides in many DNA repair proteins, including RAD51AP1. For example, S19, S21, S120, S294, and S299 in RAD51AP1 (isoform 1) were found phosphorylated upon exposure of U2OS cells to ionizing radiation (38). S294 in RAD51AP1 was found phosphorylated in HEK293 cells after UV irradiation (39). These studies suggest that S294 is a critical residue in RAD51AP1 (isoform 1) that may regulate protein activity.

We noticed that both S294 and S299 are in direct vicinity of RAD51AP1’s C-terminal bi-partite nucleic acid binding domain (9, 18). We also noticed that S294 is part of a minimal CDK consensus site (*i.e*., S/T-P) (40, 41). As such, we speculated that phosphorylation of S294 not only is required for RAD51AP1 function but may also be mediated by a CDK.

In this study, we show that RAD51AP1 bearing serine to alanine mutations at both S294 and S299 (S2A) is more proficient in binding to naked and chromatinized DNA substrates and in the stimulation of D-loop formation than wild type RAD51AP1 or RAD51AP1 with S294/S299 changed to aspartates (S2D). In cells, however, RAD51AP1-S2A confers no rescue of RAD51AP1 deficiency in toxicity tests and DNA replication assays. In contrast, RAD51AP1-S2D behaves like the wild-type protein and fully rescues RAD51AP1 deficiency. Our results support a model in which the phosphorylation of RAD51AP1-S294/299 is required for the completion of the HDR pathway in cells. We propose that RAD51AP1-S294/S299 phosphorylation is critical for RAD51AP1 release from DNA, its renewed engagement in the HDR reaction, and that S294 is a CDK2 target in early S phase cells.

## RESULTS

### RAD51AP1 cell lines for cell-based studies

Here, we studied RAD51AP1 isoform 2 (335 residues) which is the predominant isoform in human cells and the only documented species in some vertebrates (42). S294 and S299 in RAD51AP1 isoform 1 correspond to S277 and S282 in RAD51AP1 isoform 2 (Fig. 1A). Both S277 and S282 are highly conserved across different vertebrate species, as determined by ClustalW sequence alignment (Fig. 1B). We used HeLa RAD51AP1 knockout (KO) cells (43, 44) and generated clonal derivatives of RAD51AP1 KO cells that express FLAG-tagged wild type RAD51AP1 or mutants. Clonal isolates were selected to express similar levels of ectopic RAD51AP1, as determined by Western blot analysis (Figs. S1A, S1B). We selected two independently isolated clones that express RAD51AP1-S277A/S282A (S2A), two independently isolated clones that express RAD51AP1-S277D/S282D (S2D), and one clone each expressing wild type RAD51AP1 (WT) or the single residue mutants RAD51AP1-S277A or -S282A (Figs. S1A, S1B).

**Figure 1.**
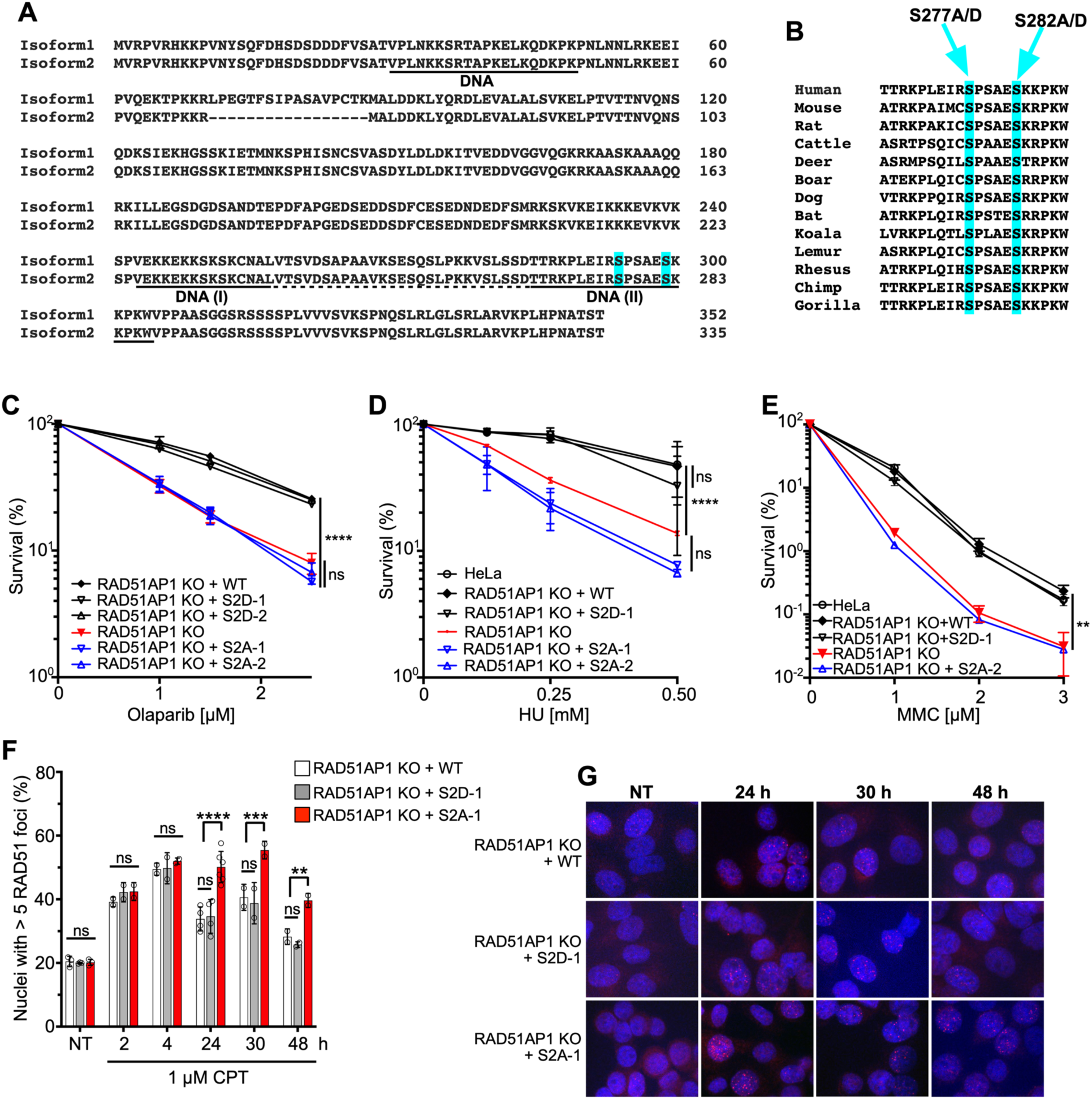
S277/282 regulate RAD51AP1 activity in HDR. *A*, ClustalW sequence homology alignment of the human RAD51AP1 isoforms 1 and 2. *B*, ClustalW homology alignment of the RAD51AP1 peptide sequences containing S277/S282 in mammals. *C*-*E*, Results from clonogenic cell survival assays to determine the sensitivity of HeLa RAD51AP1 KO cells and derivatives, as indicated (n = 3). Two-way ANOVA followed by Tukey’s multiple comparison test. \*\**p* < 0.01. \*\*\*\**p* < 0.0001. ns, not significant. *F*, Time course of RAD51 foci formation after treatment of HeLa RAD51AP1 KO cells and derivatives with 1 µM CPT for 1 h (n = 2-5). NT: not treated. Two-way ANOVA followed by Tukey’s multiple comparison test. \*\**p* < 0.01. \*\*\**p* < 0.001. \*\*\*\**p* < 0.0001. ns, not significant. *G*, Representative micrographs of RAD51 foci (red) for the results shown in (F). DAPI: blue.

To confirm that the ectopically expressed RAD51AP1 proteins retain the described RAD51AP1 interaction capabilities (8, 15, 44, 45), we performed co-immunoprecipitation experiments using anti-FLAG M2 affinity resin and cell lysates from exponentially growing cells. We show that there is no difference between FLAG-tagged RAD51AP1, RAD51AP1-S2A, and RAD51AP1-S2D in co-precipitating the previously described (8, 15, 44, 45) RAD51AP1-interacting proteins (Figs. S1C, S1D). These results suggest that the ectopically expressed RAD51AP1 proteins possess the protein interaction attributes of endogenous RAD51AP1 and likely adopt a very similar conformation.

### S277A/S282A amino acid substitutions inactivate RAD51AP1 function in cells

Next, we performed cell survival assays to test for the sensitivity of RAD51AP1-S2A and RAD51AP1-S2D cells to Olaparib, hydroxyurea (HU), and mitomycin C (MMC). We show that RAD51AP1-S2A cells are defective in rescue of RAD51AP1 deficiency in HeLa cells (Fig. 1C-1E). In contrast, RAD51AP1 or RAD51AP1-S2D fully rescue RAD51AP1 deficiency. Single mutants show partial rescue of RAD51AP1 deficiency in cell survival assays conducted with Olaparib (Fig. S1E).

Knockdown or loss of RAD51AP1 impairs the resolution of DNA damage-induced RAD51 foci (8, 9, 46). Hence, we wondered if RAD51AP1-S2A cells would be impaired in the resolution of RAD51 foci. We exposed cells to 1 µM camptothecin for 1 h and monitored the formation and resolution of RAD51 foci by immunocytochemistry. We show that RAD51AP1 KO cells with WT RAD51AP1 or RAD51AP1-S2D resolve RAD51 foci with similar kinetics; in contrast, RAD51AP1-S2A cells are significantly impaired in resolving RAD51 foci (Figs. 1F, 1G). Collectively, these results show that S277 and S282 are critical residues that regulate RAD51AP1 activity in response to DNA damage.

To understand if RAD51AP1-S2A cells are impaired in overcoming impediments to DNA replication, we used the DNA fiber assay and treated RAD51AP1 KO cells and derivatives with 4 mM HU for 5 h (Fig. 2A), a condition which essentially stalls all DNA replication (47). We determined the ability of RAD51AP1 KO cells and derivatives to restart DNA replication by measuring the speeds of the IdU tracts preceded by a CldU tract. The results show that, compared to RAD51AP1 KO cells and RAD51AP1 KO cells with WT RAD51AP1 or RAD51AP1-S2D, RAD51AP1-S2A cells are significantly impaired in the recovery from stalled DNA replication (Figs. 2B; Fig. S2A). Accordingly, RAD51AP1-S2A cells contain significantly more stalled and fewer restarted replication forks than RAD51AP1 KO, RAD51AP1 WT, and RAD51AP1-S2D cells (Fig. 2C). RAD51AP1 KO cells expressing single residue changes (S277A or S282A) also show a defect in replication fork restart that is not significantly different from that in RAD51AP1-S2A cells (Figs. S2A, S2B). Under unperturbed conditions, IdU tract replication speeds were similar in RAD51AP1 KO cells and all derivative cell lines (Figs. S2A, S2C). These results show that loss of RAD51AP1 impairs the recovery from stalled DNA replication, and that expression of mutant RAD51AP1-S2A/-S277A or - S282A further exacerbates this phenotype.

**Figure 2.**
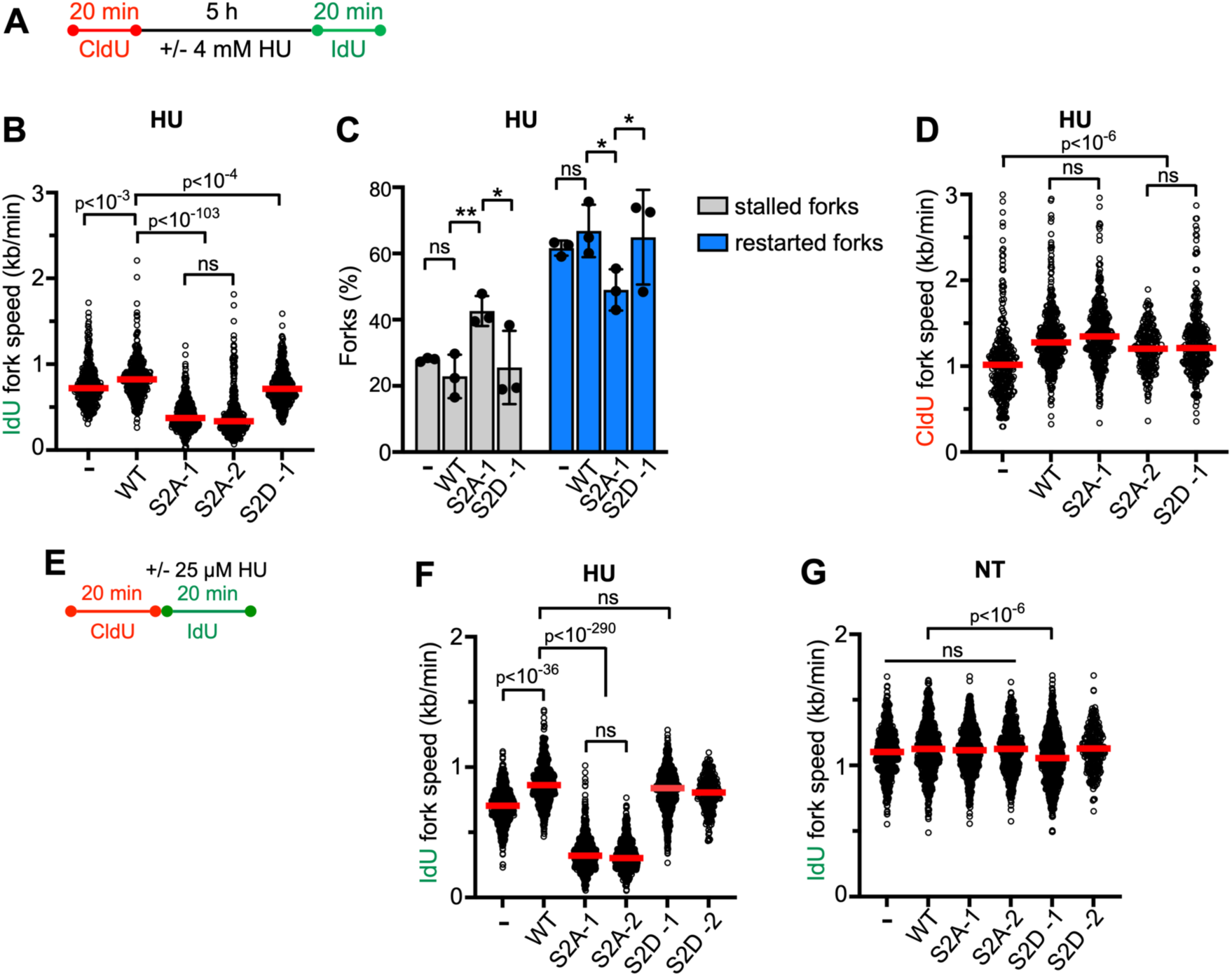
RAD51AP1-S2A cells exhibit impaired replication fork restart and fork progression in response to replication stress. *A*, Schematic of the protocol for the DNA fiber assay to determine replication restart and degradation used in (B-D). *B*, Dot plot with medians of IdU fork speeds in HeLa RAD51AP1 KO cells (−) and derivatives after HU (n = 3 for -, WT, S2A-1, S2D-1 cells; n = 2 for S2A-2 cells; 140-230 fiber tracts/experiment analyzed). *C*, Percentage of stalled and restarted forks for the results shown in (B). \**p* < 0.05. \*\**p* < 0.01. ns, not significant. *D*, Dot plot with medians of CldU fork speeds in HeLa RAD51AP1 KO cells and derivatives after HU (n = 3; 140-230 fiber tracts/experiment analyzed). *E*, Schematic of the protocol for the DNA fiber assay to determine fork progression in the presence of HU and used in (F-G). *F*, Dot plot with medians of IdU fork speeds in RAD51AP1 KO cells and derivatives treated with 25 µM HU during the IdU pulse (n = 3 for -, WT, S2A-1, S2A-2, S2D-1 cells; n = 1 for S2D-2 cells; 215-230 fiber tracts/experiment analyzed). *G*, Dot plot with medians of IdU fork speeds in unperturbed (NT) RAD51AP1 KO cells and derivatives (n = 3 for -, WT, S2A-1, S2A-2, S2D-1 cells; n = 1 for S2D-2 cells; 207-247 fiber tracts/experiment analyzed). Data were analyzed by Kruskal-Wallis test followed by Dunn’s multiple comparisons test.

To assess if RAD51AP1 and mutants protect replication forks from nuclease degradation, CldU fork speeds in cells exposed to HU were measured following the same protocol as described above (Fig. 2A). The results show that CldU fork speeds are faster in cells expressing wild type or mutant RAD51AP1 than in RAD51AP1 KO cells, and that the CldU fork speeds for all HU-treated cell lines are very similar to those in untreated cells (Figs. 2D, S2A, and S2D). These results suggest that S277 and S282 in RAD51AP1 play no major role in the protection of stalled replication forks from nuclease attrition.

Next, we monitored replication fork progression under a low concentration of HU (25 µM) given within the IdU pulse ((48); Fig. 2E). We measured IdU fork speeds in IdU tracts with a preceding CldU tract. As in the fork restart assay (Figs. 2A-C), RAD51AP1-S2A cell lines show significant defects in the fork progression assay with HU and replicate significantly slower through the IdU pulse than any of the other cell lines tested (Figs. 2F-G, S2E). In the presence of CPT (Fig. S2F), RAD51AP1-S2A cells also exhibit the most pronounced defect in fork progression (Figs. S2G-I).

### S277A/S282A enhance the association of RAD51AP1-F3 with ssDNA, dsDNA, and the NCP

We used a previously described construct and protein purification scheme to purify a C-terminal RAD51AP1 fragment (RAD51AP1-F3/F3; encompassing residues 188-335) (18, 19, 42, 49). The F3 mutant proteins (F3-S2A and F3-S2D; Figs. 3A, S3A) were overexpressed using constructs generated by site-directed mutagenesis and purified following the identical protocol as wild-type F3. RAD51AP1-F3 contains a bipartite DNA binding motif, as previously described ((18); Fig. S3B), and F3 is sufficient for RAD51AP1’s interaction with the nucleosome core particle (NCP; (19)), the minimum basic unit of chromatin in which ∼2 super-helical turns of 147 bp double-stranded DNA are wrapped around one histone octamer with no free DNA ends remaining (50, 51).

**Figure 3.**
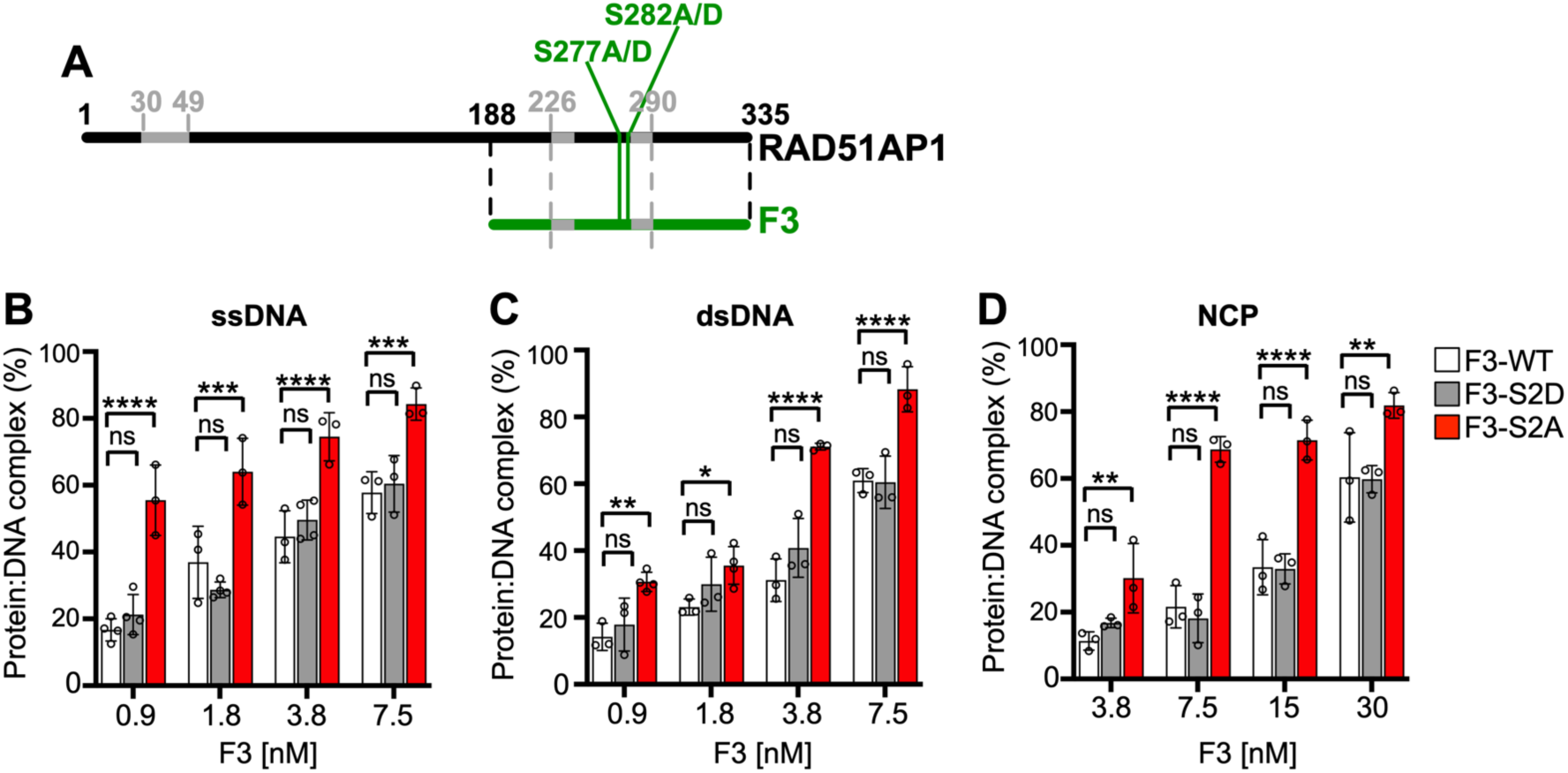
S277/282 regulate RAD51AP1 binding to ssDNA, dsDNA, and the NCP. *A*, Schematic representation of the RAD51AP1 protein (isoform 2; black) and the RAD51AP1-F3 fragment (residues 188-335; green) containing the bipartite DNA binding domain (grey) and S277 and S282. The N-terminal RAD51AP1 DNA binding domain (residues 30-49) is indicated as well. *B*-*D*, Results from EMSAs using purified MBP-F3-His_6_ and mutants with ssDNA, dsDNA, or the NCP (n = 3-4; bars represent the means ± 1 SD). Two-way ANOVA followed by Tukey’s multiple comparison test. \**p* < 0.05. \*\**p* < 0.01. \*\*\**p* < 0.001. \*\*\*\**p* < 0.0001. ns, not significant.

We tested RAD51AP1-F3 and mutants in electrophoretic mobility shift assays (EMSAs) using Cy5-labeled ssDNA, dsDNA, and the NCP (Table S1). Consistently, F3 binds all three substrates (18, 19). Notably, F3-S2A bound ssDNA, dsDNA, and the NCP more avidly than F3 or F3-S2D, while there was no significant difference between F3 and F3-S2D (Figs. 3B-D, S3C-H). Accordingly, the apparent binding constants (*K*_D(app)_) for the three different DNA substrates were lower for F3-S2A than for F3-WT and F3-S2D (Fig. S3I). These results show that S277 and S282 are critical residues engaged in regulating RAD51AP1’s affinity to both naked DNA and the NCP.

### S277A/S282A enhance RAD51 activity in duplex capture and D-loop formation

In our earlier study we showed that purified RAD51AP1 added to RAD51 filaments stimulated capturing of the NCP in the duplex capture assay (see Fig. S4A for schematic), and that the RAD51AP1-F3 domain was sufficient for the stimulation of this reaction (19). Here, we asked if the S2A mutants would show altered proficiency in duplex capture. We tested F3, F3-S2D, and F3-S2A (25-100 nM each) and show that F3-S2A is significantly more proficient in stimulating capture of the NCP than F3 or F3-S2D (Fig. S4B, S4C). In contrast, there was no significant difference between F3 and F3-S2D at all concentrations tested. Next, we purified full-length RAD51AP1 as previously described (19) and the full-length RAD51AP1-S2A and -S2D mutants (Fig. S4D). Compared to RAD51AP1 or RAD51AP1-S2D, RAD51AP1-S2A stimulated duplex capture significantly more at 50 nM (*p*<0.0001). At 100 nM, a significant difference remained between RAD51AP1 and RAD51AP1-S2A or RAD51AP1-S2D (*p*<0.01 and *p*<0.001, respectively), although RAD51AP1-S2D was as capable as RAD51AP1-S2A in stimulating the reaction (Fig. S4E, S4F; *p*=0.92).

RAD51AP1 associates with the RAD51 recombinase to stimulate joint-molecule formation (8, 9, 15, 18, 42, 49, 52), which can be assessed *in vitro* by the D-loop assay. RAD51AP1 is also capable of stimulating joint-molecule formation in the D-loop assay with chromatinized template DNA (19). Prompted by our results that show that the S2A mutant functions more efficiently in duplex capture and DNA binding, we tested full-length RAD51AP1 and mutants in the D-loop assay with both supercoiled naked and chromatinized DNA (for schematic see Fig. 4A). Addition of 100-300 nM of protein to the reaction with supercoiled DNA stimulated D-loop formation by RAD51AP1-S2A ∼1.5- to 2-fold over that of wild-type RAD51AP1 (*p*<0.0001), while RAD51AP1 and RAD51AP1-S2D show similar levels of stimulation in this reaction (Fig. 4B, 4C). In the D-loop assay with the nucleosomal array, RAD51AP1-S2A was also significantly more effective than RAD51AP1 or RAD51AP1-S2D (*p*<0.0001), although the reaction saturated for all RAD51AP1 variants at the highest concentration (300 nM) tested (Fig. 4D, 4E). As expected, the stimulation of D-loop formation by RAD51AP1 was fully dependent on RAD51 (Fig. S4G).

**Figure 4.**
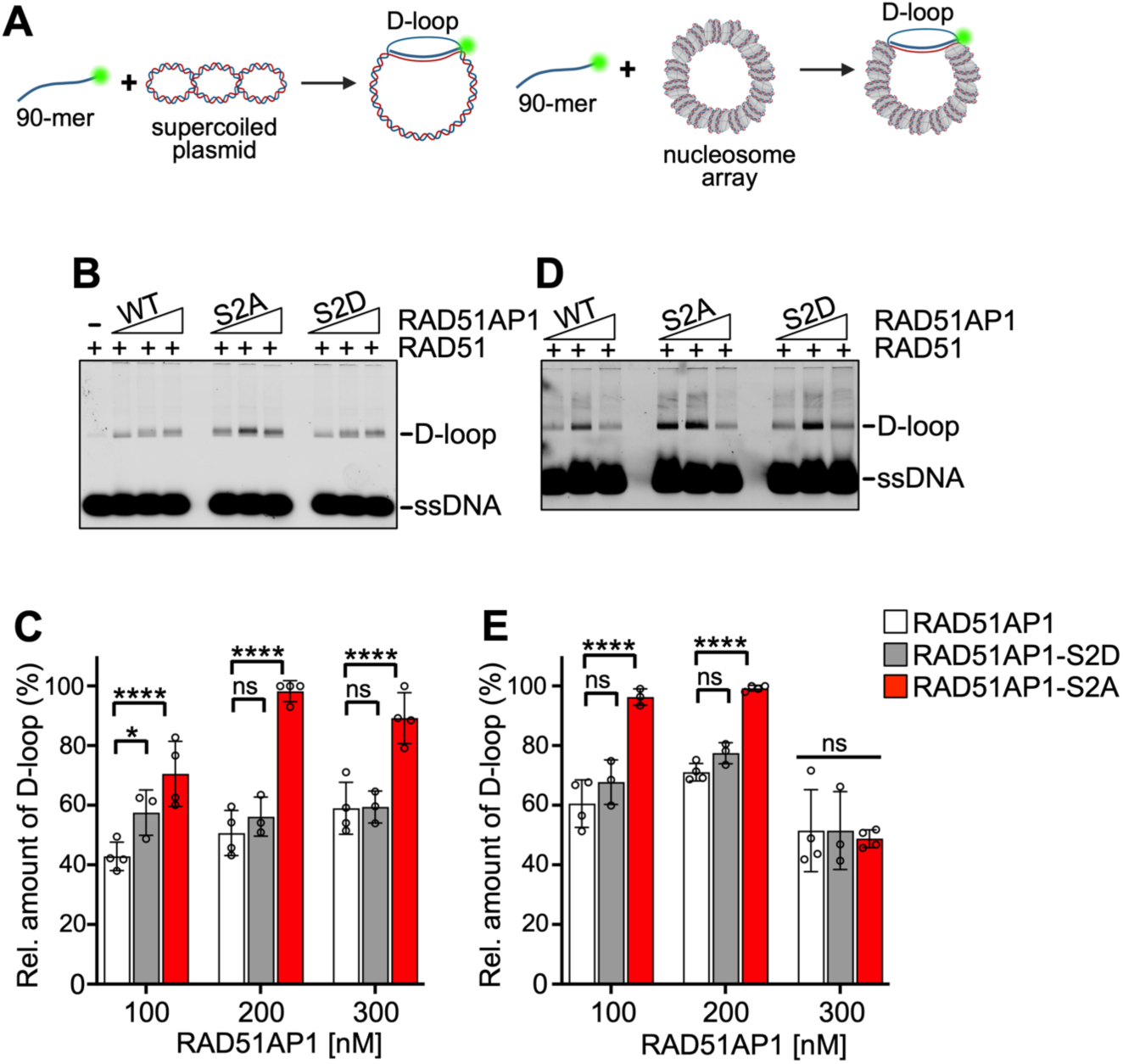
S277/282 regulate RAD51AP1’s stimulation of RAD51 in D-loop formation. *A*, Schematic of the D-loop assay with naked (left) and chromatinized (right) pBluescript II SK(−) plasmid DNA. *B*, Representative agarose gel used to quantify D-loop formation with naked DNA and His_6_-RAD51AP1-FLAG or mutants. *C*, Quantitation of D-loop formation with naked DNA (n = 3-4; bars represent the means ± 1 SD). *D*, Representative agarose gel used to quantify D-loop formation with chromatinized DNA and His_6_-RAD51AP1-FLAG or mutants. *C*, Quantitation of D-loop formation with chromatinized DNA (n = 3-4; bars represent the means ± 1 SD). Two-way ANOVA followed by Tukey’s multiple comparison test. \**p* < 0.05. \*\*\*\**p* < 0.0001. ns, not significant.

### S277 is phosphorylated by CDK1/CDK2 *in vitro*

Analysis of the primary structure of the RAD51AP1-F3 domain revealed three minimal CDK consensus phosphorylation motifs (*i.e*., S/T-P) (40, 41). S224/P225, S277/P278, and S310/P311 (Fig. 5A) also are predicted CDK sites by GPS 6.0 (53). To test if S277 indeed is a residue targeted by CDK, we purified RAD51AP1-F3, F3-S224A/S310A, and F3-S224A/S277A/S310A (Fig. S5A; for schematic see Fig. 5A) and performed *in vitro* kinase reactions with the CDK1/Cyclin B1 and CDK2/Cyclin E2 protein complexes (Fig. S5B). The results show that S277 in purified F3 is a target of both CDK2/Cyclin E2 and CDK1/Cyclin B1 (Fig. 5B, 5C; lanes 3), and that F3-S224A/S277A/S310A is not phosphorylated by either CDK complex (Fig. 5B, 5C; lanes 6).

**Figure 5.**
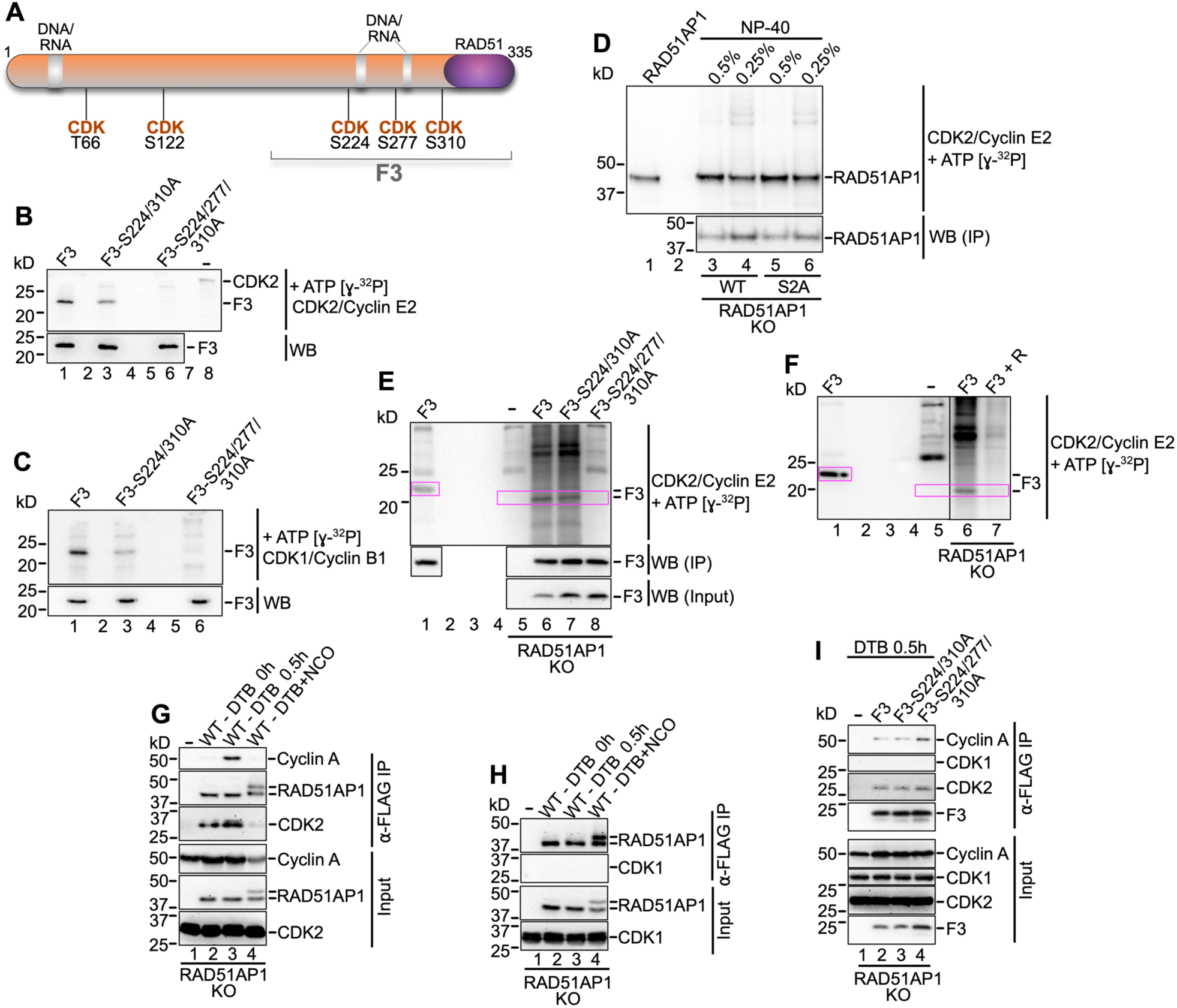
S277 is a CDK2 target in cells. *A*, Schematic of the RAD51AP1 protein (isoform 2) with predicted CDK1/2 sites (GPS 6.0). The F3 fragment (residues 188-335) and the bi-partite DNA binding domain (grey) are indicated. Predicted CDK1/2 sites in F3 (S224, S277 and S310) were changed to alanine. *B*, Results from kinase reaction of purified His_6_-F3-FLAG and mutants (S224/310A and S224/277/310A) with CDK2/Cyclin E2 and ɣ-^32^P-ATP. Upper: Phosphorimage; lower: Western blot with ⍺-FLAG antibody. *C*, Results from kinase reaction of purified His_6_-F3-FLAG and mutants (S224/310A and S224/277/310A) with CDK1/Cyclin B1 and ɣ-^32^P-ATP. Upper: Phosphorimage; lower: Western blot with ⍺-FLAG antibody. *D*, Results from kinase reaction of stably expressed (in HeLa RAD51AP1 KO cells) and precipitated FLAG-RAD51AP1 and the S2A (S277/282A) mutant with CDK2/Cyclin E2 and ɣ-^32^P-ATP. Upper: Phosphorimage; lower: Western blot with ⍺-RAD51AP1 antibody. The product of the kinase reaction with purified (from *E. coli*) His6-RAD51AP1-FLAG protein is shown for comparison purposes (lane 1). *E*, Results from kinase reaction of transiently expressed (in HeLa RAD51AP1 KO cells) and precipitated FLAG-F3 and mutants (S224/310A and S224/277/310A) with CDK2/Cyclin E2 and ɣ-^32^P-ATP. Upper: Phosphorimage; middle and lower: Western blot with ⍺-FLAG antibody. The product of the kinase reaction with purified (from *E. coli*) His_6_-F3-FLAG is shown for comparison purposes (lane 1). Pink boxes: to highlight phosphorylated F3. *F*, Results from kinase reaction of transiently expressed (in HeLa RAD51AP1 KO cells) and precipitated FLAG-F3 with CDK2/Cyclin E2 and ɣ-^32^P-ATP without and with the addition of 1 µM Roscovitine (+ R; lanes 6 and 7, respectively). The product of the kinase reaction with purified (from *E. coli*) His_6_-F3-FLAG is shown for comparison purposes (lane 1). *G*, CDK2 and Cyclin A are present in ⍺-FLAG precipitates of FLAG-RAD51AP1 (WT) stably expressed in HeLa RAD51AP1 KO cells, synchronized by a double thymidine block (DTB) and released into early S phase (lane 3). CDK2 and Cyclin A are not present FLAG-RAD51AP1 (WT) precipitates from cells arrested in late G2/M phase by treatment with nocodazole (NCO; lane 4). Note: In G2/M phase cells (lane 4), a partial mobility shift is observed for RAD51AP1 that – based on our preliminary results – is likely to be unrelated to phosphorylation (data not shown). *H*, CDK1 is not present in FLAG-RAD51AP1 (WT) precipitates of cells synchronized in early S (lane 3) or late G2/M phase (lane 4). *I*, CDK2 and Cyclin A are present in precipitates of transiently expressed FLAG-F3 and mutants in RAD51AP1 KO cells synchronized by a DTB and released into early S phase (lanes 2-4).

### S277 is a target of CDK2 in early S phase cells

To interrogate the CDK specificity for RAD51AP1-S277 phosphorylation within the context of cells, we used the previously described HeLa RAD51AP1 KO cell lines that stably express FLAG-RAD51AP1 or FLAG-RAD51AP1-S2A, precipitated the ectopic protein with anti-FLAG M2 affinity resin, and subjected the eluted proteins to *in vitro* kinase reactions with the CDK2/Cyclin E2 or CDK1/Cyclin B1 protein complexes. Both RAD51AP1 and RAD51AP1-S2A were readily phosphorylated by CDK2/Cyclin E2 (Fig. 5D) but were resistant to phosphorylation by CDK1/Cyclin B1 (data not shown). No difference was detected in the strengths of the phospho-signals between RAD51AP1 and RAD51AP1-S2A after CDK2/Cyclin E2 phosphorylation (Fig. 5D), likely due to the prevalence of several predicted CDK2 sites within full-length RAD51AP1 (GPS 6.0 (53); Fig. 5A).

We then reverted to transiently expressing the FLAG-tagged F3 fragment and F3 mutants (S224A/S310A and S224A/S277A/S310A) in RAD51AP1 KO cells. Anti-FLAG M2 affinity resin was used to purify the F3 variants, and the eluted proteins were subjected to *in vitro* kinase reactions. F3 and F3-S224/310A are phosphorylated by CDK2/Cyclin E2, as shown in Fig. 5E (lanes 6 and 7, respectively). Mutagenesis of S277 to alanine, abrogates CDK2/Cyclin E2 phosphorylation (Fig. 5E, lane 8). Addition of 1 µM Roscovitine, a selective CDK inhibitor, to the reaction also abrogates F3 phosphorylation by CDK2/Cyclin E2 (Fig. 5F, lane 7). However, *in vitro* phosphorylation with CDK1/Cyclin B1 of F3, F3-S224/310A, and F3-S224/277/310A led to a faint phospho-signal only with no progressive loss of signal for the mutants (Fig. S5C, lanes 6-8). These results suggest that RAD51AP1-S277 likely is not a major target of CDK1.

Next, we synchronized RAD51AP1 KO cells expressing FLAG-RAD51AP1 by a double thymidine block (DTB; (54)) and assessed the interactions between RAD51AP1 and CDK1 or CDK2 in anti-FLAG precipitates with or without treatment of cells with nocodazole. CDK2 and Cyclin A were present in anti-FLAG precipitates from early S phase cells (Fig.S5D, Fig. 5G, lane 3). As expected, neither CDK2 nor Cyclin A were present in anti-FLAG precipitates from G2/M phase cells after the cells were released from the DTB into medium containing nocodazole (Fig. S5D, Fig. 5G, lane 4). There was no evidence of an interaction between RAD51AP1 and CDK1 at either cell cycle stage (Fig. 5H). Transiently expressed F3 and the F3 mutants retain the ability to interact with CDK2 and Cyclin A in early S phase cells (Fig. 5I, lanes 2-4). Collectively, these results strongly suggest that RAD51AP1-S277 is a target of CDK2 in early S phase cells.

## DISCUSSION

RAD51AP1 directly interacts with RAD51 and is an established enhancer of RAD51 activity in HDR. Yet, the mechanistic details of how RAD51AP1 is controlled throughout the individual stages of the HDR reaction have remained largely obscure. Several quantitative phospho-proteomic studies have provided indirect evidence of RAD51AP1 regulation by phosphorylation (38, 39, 55, 56). There has been no report, however, to directly test and further interrogate the role of any of the many phosphorylated residues in RAD51AP1.

Here, we concentrated on S277 and S282, two serines phosphorylated in human RAD51AP1 (isoform 2) upon exposure of cells to DNA damage (38, 39). We used genetic and biochemical approaches to directly assess the consequences of S277/S282 phosphorylation for RAD51AP1 function in cells and provide the first direct evidence of RAD51AP1 regulation by phosphorylation. We show that phosphorylation of both S277 and S282 are critical for RAD51AP1 function and cell survival. Moreover, our results are in support of S277 being a target of CDK2, linking active RAD51AP1 to cell cycle stage.

CDKs regulate cell cycle progression and target many components of the DNA damage response (57). CDKs directly modulate proteins in DSB repair, including proteins that function in the HDR pathway (29, 36, 58–63). Our results expand this list of proteins to include RAD51AP1.

Compared to wild type RAD51AP1 and RAD51AP1-S277/282D, RAD51AP1-S277/282A displays greater proficiency in DNA binding, duplex capture, and D-loop formation. Yet, RAD51AP1-S277/282A is entirely defective in rescuing RAD51AP1 deficiency in cells. Of note, RAD51AP1-S277/282A retains the reported RAD51AP1 protein interactions (8, 15, 44, 45), suggesting that this mutant adopts a conformation similar to the wild-type protein in cells. Based on these results, we propose that RAD51AP1-S277/282A is defective in facilitating the required hierarchy of individual steps in HDR (*i.e*., pre-synapsis, synapsis, and strand invasion; Fig. 6), that involve RAD51AP1 engagement (8, 9, 14). We suggest that loss of S277/282 phosphorylation limits RAD51AP1 protein flexibility and the dynamics that are required to productively move the HDR reaction forward.

**Figure 6.**
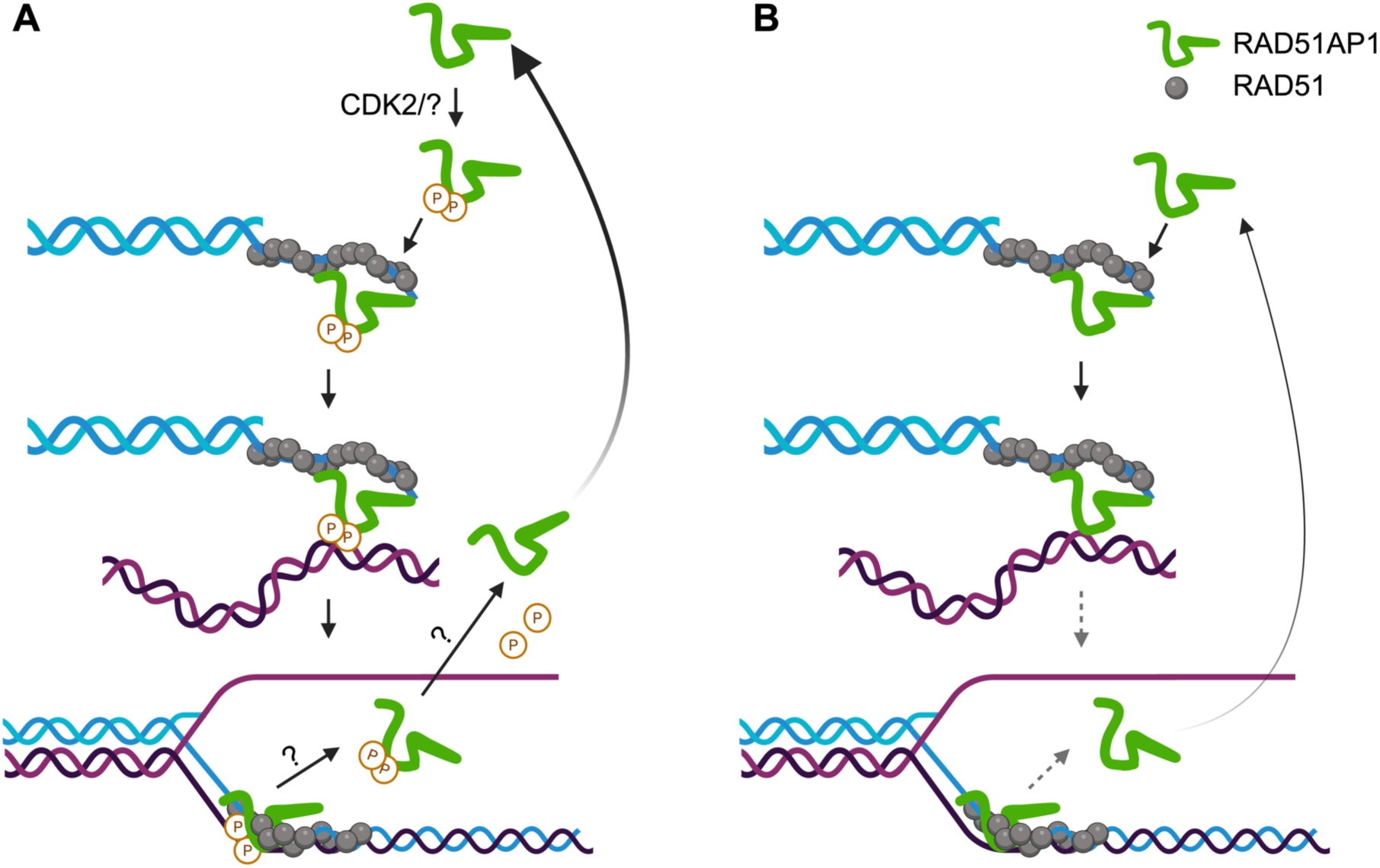
Model to illustrate how RAD51AP1-S277/S282 phosphorylation may lead to RAD51 activation in synapsis and D-loop formation. *A*, S277/S282 phosphorylation may occur prior to RAD51AP1’s association with the presynaptic filament. Alternatively, S277/S282 phosphorylation may occur when RAD51AP1 is bound to the presynaptic filament. Phosphorylation may allow RAD51AP1 to engage transiently and hierarchically in synapsis and strand exchange. It is unclear how RAD51AP1 leaves the D-loop. RAD51AP1 is dephosphorylated prior to its renewed engagement in HDR and when CDK2 is active. The kinase that targets S282 and the phosphatases involved remain to be identified. *B*, The RAD51AP1-S277/282A mutant displays enhanced proficiency in DNA binding, synapsis and D-loop formation, but is defective in promoting the continuous reaction due to its lack in flexibility and compromised dynamics. An HDR defective phenotype is observed. Created with BioRender.com.

S277/282 are embedded in RAD51AP1’s bi-partite DNA binding domain, a domain that is sufficient for its association with nucleosomes and predicted to play a major role in the formation of a molecular bridge between the molecules undergoing exchange (9, 18, 19). Although the molecular details that initiate the release of RAD51AP1 after D-loop formation remain unclear, we suggest that S277/282 phosphorylation may persist during RAD51AP1 disengagement (Fig. 6), and that RAD51AP1 release may be triggered by additional modification(s). One such modification could be at S310 (S327 in human RAD51AP1 isoform 1), previously identified as both a NEK2 and CDK1 target in mouse embryonic fibroblasts (55).

RAD51AP1 is an intrinsically disordered protein (IDP) (64). IDPs are characterized by their lack of a well-defined structure, and they are predicted to be highly dynamic and flexible (65). Moreover, bioinformatic studies have shown that phosphorylation of IDPs and their intrinsically disordered regions (IDRs) occurs more frequently than phosphorylation of folded proteins (65, 66). PONDR (pondr.com) identifies four IDRs in RAD51AP1, the longest of which encompasses the 91 most C-terminal amino acids. This IDR contains RAD51AP1’s bi-partite DNA binding domain, S277/282, and 13 additional residues that can undergo phosphorylation (phosphosite.org). Given such complexity, we are aware that our model presented in Fig. 6 is oversimplified and incomplete. In this regard, the RAD51AP1 mobility shift that we observed in G2/M phase cells by Western blot and that may be unrelated to phosphorylation (data not shown), also indicates a more involved cell cycle-specific regulation of the RAD51AP1 protein.

In cell survival assays, we obtained an intermediate phenotype for cells expressing RAD51AP1 with a single residue change (S277A or S282A). These results suggest that phosphorylation of both S277 and S282 is important for proper regulation of RAD51AP1 function. Using the DNA fiber assay to monitor the short-term response of the RAD51AP1 variants to replication stress, we find that the RAD51AP1 single mutants are as impaired in replication restart or fork progression as the RAD51AP1-S277/282A double mutant. Collectively, these results suggest that compensatory mechanisms may counteract the lack of the second phosphorylated serine in longer-term cell survival studies.

One limitation of our study is that we have not elucidated the possibly complex functional interaction between S277 and S282 phosphorylation. As such, it remains unclear if the S277/282 phosphorylations are dependent, independent, cooperative, or underlie a clear-cut hierarchy (possibly associated with distinct stages of RAD51 stimulation). From our results in cells, we infer that multi-site phosphorylation widens RAD51AP1 function, making complex regulatory mechanisms possible, including hierarchical and co-operative events. Based on the two positively charged lysines immediately downstream of S282, S282 may be a target of a basophilic kinase, such as PKA or PKC, both of which were shown to have roles in the DNA damage response. For example, sequential phosphorylation of p19INK4d by CDK2 (S76) and PKA (T141) is critical for cell survival after genotoxic stress (67), and phosphorylation by PKC of HMGB1-S46, a protein that binds to damaged DNA and activates proteins in nucleotide and base excision repair (68), significantly reduces its binding affinity to DNA (69). Future investigations will need to be directed towards identifying the kinase that targets S282, the functional interaction between S277 and S282 phosphorylation, and the detailed regulatory mechanisms of RAD51AP1 activation.

RAD51AP1 is an attractive future target in cancer therapy. This is because elevated RAD51AP1 is associated with therapy resistance and poor prognosis for several different cancer types (14, 25, 26, 70, 71). Moreover, RAD51AP1 plays a critical role in the alternative lengthening of telomeres (ALT) pathway (22, 72, 73), a pathway that is used by ∼10-15% of cancers in cancers and that typically grow aggressively (74). As such, a better understanding of the molecular details of RAD51AP1 regulation, as we have begun to uncover here, is critical and has the potential to guide the design of new and improved cancer therapeutics.

## EXPERIMENTAL PROCEDURES

### Cell lines, Site-Directed Mutagenesis, and Western Blots

HeLa cells were obtained from ATCC and maintained as recommended. HeLa cells that are knockout (KO) for *RAD51AP1* were described previously (43, 44). RAD51AP1 KO cells were used to generate RAD51AP1-wild-type (-WT), RAD51AP1-Ser277/282Ala (-S2A) and RAD51AP1-Ser277/2822Asp (-S2D) cells. First, RAD51AP1 was amplified from pOK24 (15) using the primer pairs listed in Table S1. Then, the PCR product was cloned from *Sal*I to *Xho*I into pENTR1A (Thermo Fisher Scientific). Mutations in RAD51AP1 were generated in pENTR1A-FLAG-RAD51AP1 using the primers listed in Table S1 and Q5 Site-Directed Mutagenesis Kit (New England Biolabs). pENTR1A constructs were transferred into pLentiCMV/TO DEST#2 (75) using Gateway LR Clonase II (Thermo Fisher Scientific) for the production of lentiviral particles in HEK293FT cells (Thermo Fisher Scientific), as described (75). Lentivirus was used to transduce *RAD51AP1 KO* cells in 6 µg/ml polybrene, as described (75), and clonal isolates with similar expression levels were selected in 2 mg/ml G418.

For the transient expression of RAD51AP1-F3, -F3-S224/310A, and -F3-S224/277/310A in RAD51AP1 KO cells, F3, F3-S224/310A, and F3-S224/277/310A were amplified from the respective bacterial expression plasmids (in pQE-80L) using the primer pairs listed in Table S1. The PCR products were cloned from *Not*I to *Bam*HI into pcDNA3.1(−), and transient transfections were performed as described previously (76).

Western blot analyses were performed following our standard protocols (77). The primary antibodies that were used are: α-RAD51AP1 ((52); 11255-1-AP; ProteinTech; 1:10,000); α-RAD51 (Ab-1; EMD Millipore; 1:3,000); α-PARP1 (ab6079; Abcam; 1:1,000); α-MSH2 (ab52266; Abcam; 1:10,000); α-Tubulin (DM1A; Santa Cruz Biotechnology; 1:1,000); α-FLAG (F3165; Sigma; 1:1,000); α-MBP (PAI-989; Thermo Fisher Scientific; 1:5,000); α-RAD54L (F-11; sc-374598; Santa Cruz Biotechnology; 1:500); α-NUCKS1 ((78); 1:10,000); α-GST (ab19256; Abcam; 1:10,000); α-UAF1 (ab122473; Abcam; 1:100); α-CDK1 (10762-1-AP; ProteinTech; 1:2,000); α-CDK2 (10122-1-AP; ProteinTech; 1:20,000); α-Cyclin A (H-432; Santa Cruz Biotechnology; 1:1,000). HRP-conjugated goat anti-rabbit or goat anti-mouse IgG (Jackson ImmunoResearch; 1:10,000) were used as secondaries. Western blot signals were acquired using a Chemidoc XRS+ gel imaging system and ImageLab software version 5.2.1 (BioRad).

### Cell Survival Assays

Clonogenic cell survival assays after treatment of cells with Mitomycin C (MMC; Sigma) or in the presence of Olaparib (AZD2281; Selleck Chemicals) were performed as described (44). To assess sensitivity of cells to hydroxyurea (HU; Sigma), a colorimetric cell proliferation assay with sulforhodamine B (SRB) was used, as described (79). Briefly, 1000 cells/well were plated in triplicate in regular growth medium in a 96-well plate. After 24 h, HU was added at the concentrations as specified, and cells were incubated for 4 days. Cells were washed in warm PBS three times, fresh medium was added, and cells were incubated for 2 days. Without removing the medium (200 µl/well), cells were fixed by the addition of 100 µl 10% trichloroacetic acid at 4°C for 1 h. Plates were washed with tap water four times and air dried. Cells were stained in 0.05% SRB solution (in 1% glacial acetic acid) with gentle shaking at room temperature for 30 min. Then, plates were washed three time with 200 µl 1% glacial acetic acid/well and air dried. SRB was solubilized in 10 mM Tris-HCl, pH 8.0, and optical densities were determined at OD560 nm using a Modulus microplate reader (Turner Biosystems). Densities were normalized to cells not treated with HU to assess cellular sensitivities.

### Co-immunoprecipitations

HeLa and RAD51AP1 KO cells expressing wild type RAD51AP1 or RAD51AP1 mutants (∼6×10^6^ cells each) were trypsinized, lysed in ice-cold lysis buffer (50 mM Tris-HCl, pH 7.5, 300 mM NaCl and 0.5% NP40 containing EDTA-free Protease (Roche) and HALT Phosphatase inhibitors (Thermo Fisher Scientific). Cell lysates were cleared by centrifugation at 10,000 × g and diluted to 25 mM Tris-HCL, pH 7.5, 150 mM NaCl, 0.1% NP40 containing 0.1 U DNaseI/µg (Gold Biotechnology) protein. Cell lysate containing 2 mg total protein was incubated with 25 µl anti-Flag M2 resin (Sigma) at 4°C for 90 min. Trapped complexes were washed three times in 250 µl binding buffer, and bound proteins were eluted in 40 µl binding buffer containing 150 ng/ µl/ 3×Flag peptide (Sigma). Eluted protein complexes were mixed with 8 µl 6× Laemmli buffer (Thermo Fisher Scientific), fractionated on Tris-glycine gels and subjected to western blot analysis.

### DNA Fiber Assay

The DNA fiber assay was performed as described (44, 80). The fibers were spread, imaged, and measured as described (80, 81). Two slides per sample were prepared for each experimental repeat, and each pair of slides was blinded after immunodetection to avoid bias. Fiber tracts were converted to kb, as described (48), using a conversion factor of 2.59 kb/µm (82). Fork speeds (kb/min) were obtained by dividing the lengths of the tracts (in kb) by the labeling time (in min).

### Cell Synchronization, Flow Cytometry, and Cell Cycle Analysis

Flow cytometry, the double-thymidine block (DTB), and cell cycle analyses were carried out as described (44, 76). For treatment with Nocodazole (NCO; AdipoGen Life Sciences), cells were released into regular growth medium for 4 h after the DTB before NCO (80 ng/ml) was added, and cells were incubated for another 14 h before cell harvest and lysis.

### DNA Substrates, Nucleosome Core particles (NCPs), Chromatin Assembly, Electromobility Shift Assays (EMSAs)

Biotinylated and Cy5-labeled ssDNA oligos (Table S1) were obtained from IDT. A Cy5-labeled 147 bp dsDNA fragment was generated by PCR amplification from pUC19 plasmid DNA containing a single copy of the 147 bp 601 Widom fragment (83), using the primer pair listed in Table S1. The PCR product was purified by FPLC (Pharmacia Biotech) using a MonoQ 5/50 GL column. The bound DNA fragment was eluted using a linear 0-1.0 M NaCl gradient and eluted at 400 mM NaCl. Peak fractions were pooled and precipitated in 0.1 % (vol/vol) 3 M sodium acetate and 3 × vol 100% ice-cold ethanol. The pellet was washed once with 70% ethanol, air dried, resuspended in TE, pH 8.0, and stored in −20°C until further use.

NCPs were reconstituted using the purified 5’-Cy5-labelled 147 bp dsDNA fragment with positioning sequence (601 Widom fragment) and human histone octamer by serial salt dilution as described (19, 50, 84, 85). NCP quality was assessed by native 5% polyacrylamide gel electrophoresis (PAGE) as described (19, 76). Chromatin was assembled with human histone octamer by salt gradient dialysis on pBluescript II SK(−) plasmid DNA as described (86). The plasmid/octamer ratio was based on 207 ± 4 bp DNA/nucleosome. The quality of the assembled chromatin was controlled by limited digestion with MNase.

EMSAs were performed with some modifications as described (19). Briefly, Cy5-labeled DNA or NCP (10 and 40 nM, respectively) were incubated with 0.315-60 nM wild type RAD51AP1, RAD51AP1-F3 or mutants in buffer A (50 mM Tris-HCl, pH 7.5, 100 mM NaCl, 2 mM DTT) at 4°C for 30 min. DNA/protein or NCP/protein complexes were fractionated by native 5% PAGE gels in 0.2× TBE at 150 V for 60 min. The D-loop assay was performed as described (76, 87). Image acquisition occurred on a Typhoon biomolecular imager (Cytiva), and signals were quantitated by ImageJ software.

### Purification of RAD51AP1, RAD51AP1 Fragments and Mutants

The expression constructs for His_6_-RAD51AP1-FLAG and MBP-RAD51AP1-F3-His_6_ are described elsewhere (18, 76). To generate the construct for His_6_-F3-FLAG the RAD51AP1-F3 fragment was PCR amplified from His_6_-RAD51AP1-FLAG using the primer pairs listed in Table S1 and cloned form *Bam*H1 to *Sal*I into pQE-80L (Qiagen). Mutations in RAD51AP1 or in RAD51AP1-F3 were introduced using the primers listed in Table S1 and Q5 Site-Directed Mutagenesis (New England Biolabs). RAD51 was expressed and purified as described (88, 89). Purified human histone octamer was obtained from the Histone Source at Colorado State University (https://www.histonesource.com/) and purified as described (90). His_6_-RAD51AP1-FLAG, His_6_-F3-FLAG, and MBP-F3-His_6_ were purified as described (19, 76).

### Pull-downs

FLAG pull-downs were performed using anti-FLAG M2 affinity resin (Sigma), essentially as described by the manufacturer. Briefly, anti-FLAG resin was equilibrated in binding buffer (50 mM Tris-HCl at pH7.5, 150 mM NaCl, 0.1% Triton X-100, and 100 µg/ml BSA). Purified RAD51AP1 or RAD51AP1-F3 (100 nM each) was added to the equilibrated beads and incubated at 4°C for 1 h with gentle agitation. Unbound protein was removed from the resin by centrifugation at 1,500 × g for 3 min. Recombinant CDK1/Cyclin B1 or CDK2/Cyclin E2 (Sigma; 150 nM) was added in binding buffer and incubated at 4°C for 2 h with gentle agitation in the presence of DNase I (1 U/µg protein). The resin was washed twice in 200 µl binding buffer, and bound protein complex was eluted in 40 ul elution buffer containing150 ng/µl 3 × FLAG peptide. Eight microliters of 6 × Laemmli buffer were added, and samples were fractionated by 10-15% SDS-PAGE for Western blot analysis.

### Kinase Assays

To phosphorylate purified wild type RAD51AP1, RAD51AP1-F3 or mutants a standard kinase reaction using ɣ-^32^P-ATP (6000 Ci/mmol) was performed, as described (91). Briefly, 50 ng of the purified proteins were incubated with 25 ng CDK1/Cyclin B1 or CDK2/Cyclin E2 (Sigma) in kinase buffer (12.5 mM Tris-HCl, pH 7.5, 0.2 mM EDTA, 10 mM magnesium acetate, 1 mM DTT). The reaction was initiated by the addition of 2.5 µCi ɣ-^32^P-ATP and incubated at 30°C for 30 min. Then, 1.33 µM cold ATP was added and incubated for another 5 min for the completion of reaction. The reaction was stopped by the addition of 6× Laemmli buffer. The protein was fractionated on 10 or 15% SDS-PAGE and transferred to a PDVF membrane. Phosphor-imaging was performed using a Typhoon biomolecular imager (Cytiva).

For the phosphorylation of immunoprecipitated RAD51AP1, RAD51AP1-F3, and mutants from HeLa cells, 2×10^6^ cells were seeded in regular growth medium and precipitations with anti-FLAG M2 affinity resin (Sigma) were carried out as described above. Eluted RAD51AP1/F3 was stored on ice overnight and used as substrate in kinase reactions with CDK1/Cyclin B1 or CDK2/Cyclin E2 (Sigma), as described above.

### Statistical analyses

Statistical analyses were performed using Prism 10 GraphPad Software on the data from 2-4 independent experiments using two-way ANOVA followed by Tukey’s multiple comparison test or Kruskal-Wallis test followed by Dunn’s multiple comparisons test *P* ≤ 0.05 was considered significant.

## Data availability

All data are included in the manuscript and the supporting information file. The raw data are available upon request.

## Author contributions

N.S., M.U. data curation, data analyses, validation; Y.K. and P.S. resources; C.W., N.S. conceptualization; C.W. funding acquisition, writing of the manuscript, project administration.

## Funding and additional information

This work was supported by National Institutes of Health Grants R01GM144579 (C.W.), R50CA265315 (Y.K.), and R01ES007061 (P.S.).

The content is solely the responsibility of the authors and does not necessarily represent the official views of the National Institutes of Health.

## Abbreviations

The abbreviations used are:

AA: Amino acid
ANOVA: Analysis of variance
ALT: Alternative lengthening of telomeres
ATP: Adenosine triphosphate
BSA: Bovine serum albumin
CDK: Cyclin-Dependent Kinase
CtIP: C-terminal binding protein-Interacting Protein
CPT: Camptothecin
D-loop: Displacement-loop
DR-loop: DNA/RNA-loop
EMSA: Electrophoretic Mobility Shift Assay
HPLC: High-Pressure Liquid Chromatography
HDR: Homology-Directed Repair
HU: Hydroxy Urea
ICLs: Inter-strand Crosslinks
MMC: Mitomycin C
NCO: Nocodazole
MNase: Micrococcal Nuclease
MBP: Maltose-Binding Protein
MNase: Micrococcal Nuclease
NCP: Nucleosome Core Particle
Ni-NTA: Nickel Nitrilotriacetic Acid
PIN1: Peptidyl-Prolyl Cis-Trans Isomerase NIMA-Interacting 1
PVDF: Polyvinylidene Difluoride
UAF1: USP1-Associated Factor 1
RAD51AP1: RAD51-Associated Protein 1
RIF1: Replication Timing Regulatory Factor 1
RPA: Replication Protein A
UHRF1: Ubiquitin-like, containing PHD and RING finger domains 1
PKA: Protein Kinase A
PKC: Protein Kinase C
HMGB1: High-mobility Group Box 1
UV: ultraviolet
d.f.: degrees of freedom
r^2^: coefficient of determination

