## Supporting Information for "Phosphoregulation of RAD51AP1 function in homology-directed repair"

**Supporting Figure S1**
**Supporting Figure S2**
**Supporting Figure S3**
**Supporting Figure S4**
**Supporting Figure S5**

**Supporting Table S1**

**Supporting References**

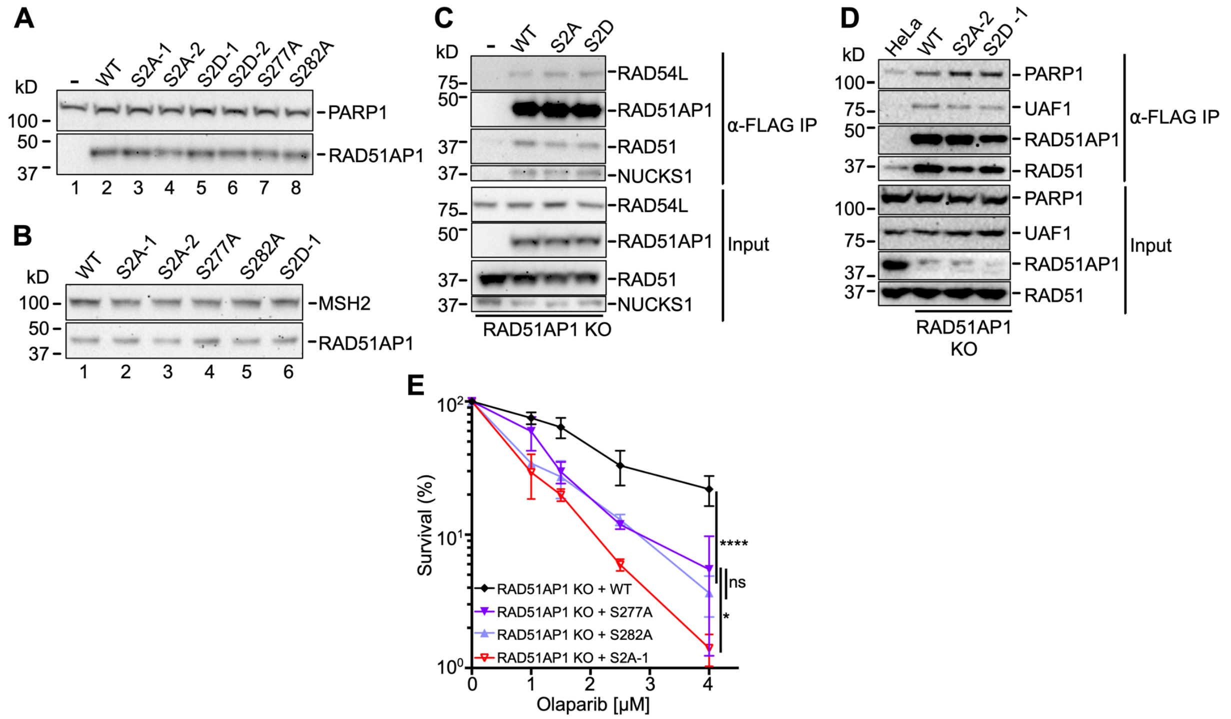


**Figure S1.** Related to Figure1. **RAD51AP1 variants retain the known RAD51AP1 protein interactions in cells.** *A-B*, Western blots to show the expression of ectopic RAD51AP1 and mutants in HeLa RAD51AP1 KO cells. For RAD51AP1-S2A and RAD51AP1-S2D, two independently isolated clones were analyzed. PARP1, MSH2: loading controls. *C-D*, Western blots after anti-FLAG co-immunoprecipitations to show that ectopic RAD51AP1 and RAD51AP1 mutants (-S2A and -S2D) retain the described protein interactions of endogenous RAD51AP1(1-4). *E*, Results from clonogenic cell survival assays in response to Olaparib to determine the sensitivity of cells expressing RAD51AP1 with single residue changes (S277A or S282A) in comparison to cells expressing RAD51AP1-S2A. Two-way ANOVA followed by Tukey's multiple comparison test. *$p < 0.05$. ****$p < 0.0001$. ns, not significant.

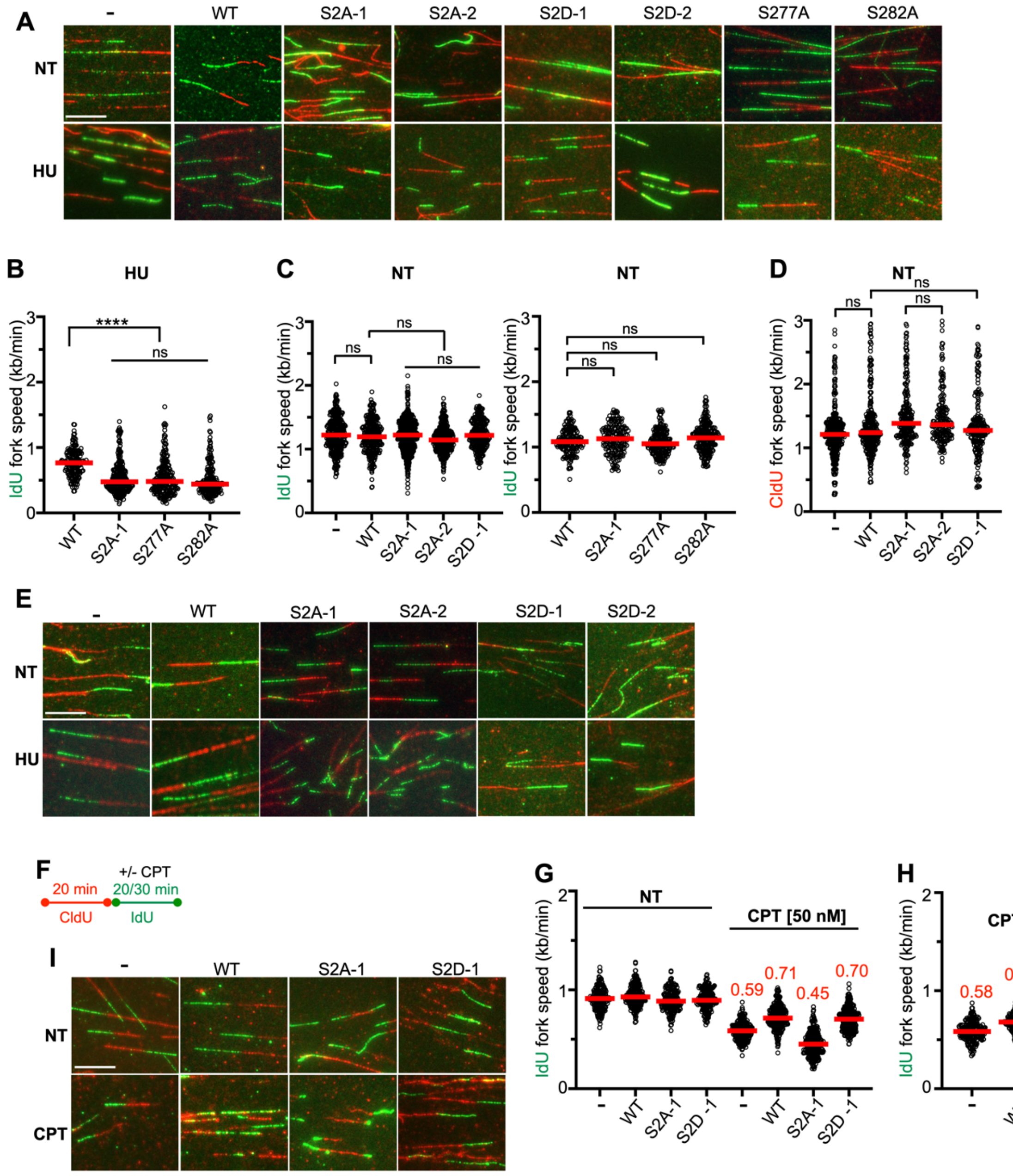


**Figure S2.** Related to Figure 2. **RAD51AP1-S2A, -S277A, and -S282A cells exhibit impaired replication fork restart and fork progression in response to replication stress.** *A*, Representative micrographs of DNA fibers in HeLa RAD51AP1 KO cells and derivatives under unperturbed conditions (NT) and after HU to determine recovery from stalled replication by IdU fork progression (green; Fig. 2A-C) and fork resection by CldU fork progression (red; Fig. 2A, D). *B*, Dot plot with medians of IdU fork speeds in HeLa RAD51AP1 KO cells expressing RAD51AP1 or mutants to show fork restart after HU (labeling protocol shown in Fig. 2A; n = 3;100-150 fiber tracts/experiment analyzed). *C*, Dot plot with medians of IdU fork speeds in unperturbed HeLa RAD51AP1 KO cells (-) and derivatives (n=3; 100-150 fiber tracts/experiment analyzed). *D*, Dot plot with medians of CldU fork speeds in unperturbed HeLa RAD51AP1 KO cells (-) and derivatives (n=3 for -, WT, S2A-1, S2D-1 cells; n = 2 for S2A-2 cells; 100-150 fiber tracts/experiment analyzed). Data were analyzed by Kruskal-Wallis test followed by Dunn's multiple comparisons test. ****$p$ < 0.0001. ns, not significant. *E*, Representative micrographs of DNA fibers in HeLa RAD51AP1 KO cells (-) and derivatives under unperturbed conditions (NT)

and after mild replication stress (labeling protocol shown in Fig. 2E). *F*, Schematic for the protocol of the DNA fiber assay to determine fork progression in the presence of mild replication stress by camptothecin (CPT) and used in (G-I). *G*, Dot plot with medians of IdU fork speeds in HeLa RAD51AP1 KO cells (-) and derivatives without (NT) and in the presence of 50 nM CPT (n = 1; 150-200 fiber tracts analyzed). *H*, Dot plot with medians of IdU fork speeds in HeLa RAD51AP1 KO cells (-) and derivatives in the presence of 25 nM CPT (n = 1; 150-200 fiber tracts analyzed). *I*, Representative micrographs of DNA fibers in HeLa RAD51AP1 KO cells (-) and derivatives under unperturbed conditions (NT) and after mild replication stress (25 nM CPT). Scale bars: 10 µm.

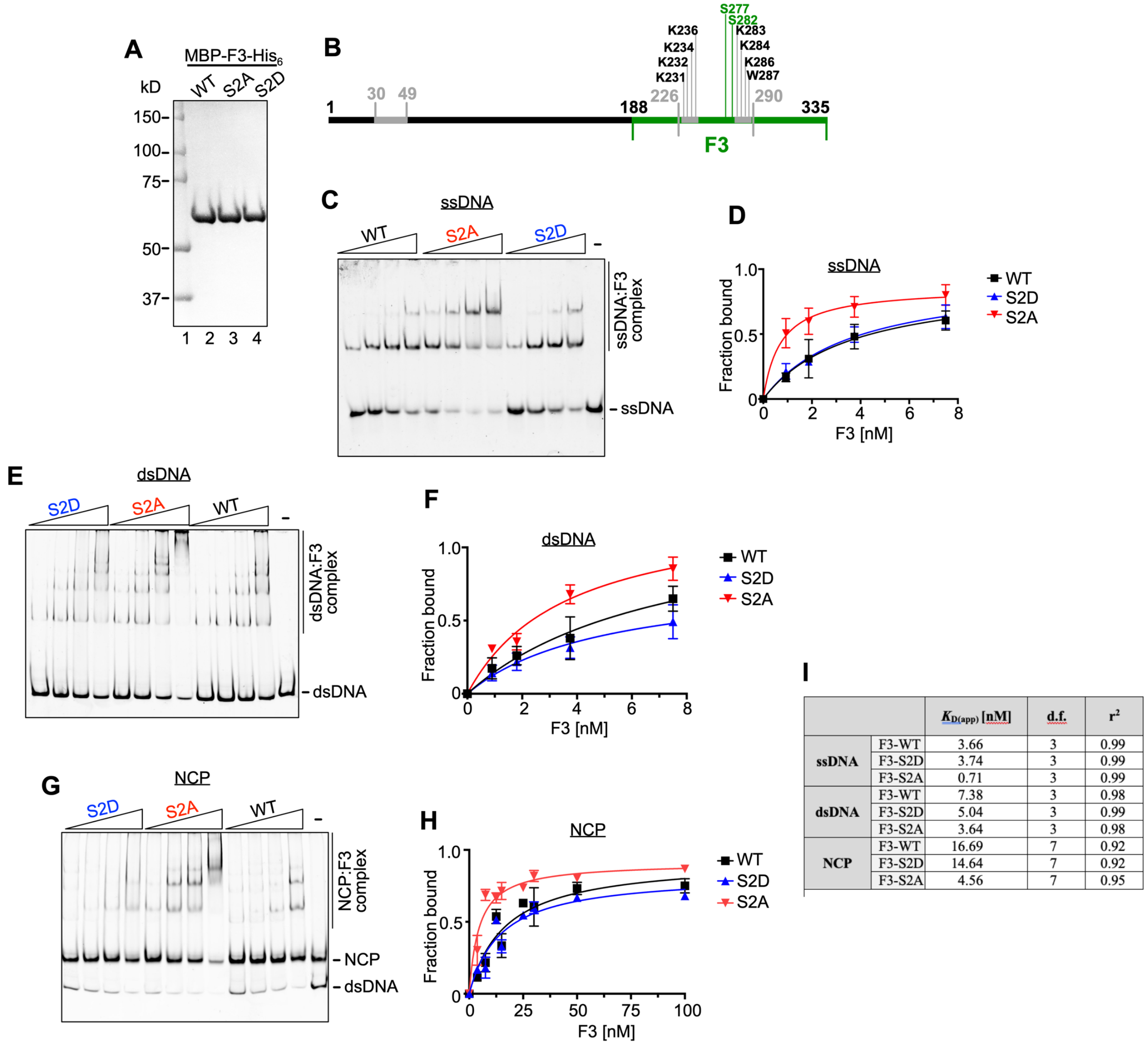


| | | $K_{D(app)}$ [nM] | d.f. | $r^2$ |
|---|---|---|---|---|
| ssDNA | F3-WT | 3.66 | 3 | 0.99 |
| | F3-S2D | 3.74 | 3 | 0.99 |
| | F3-S2A | 0.71 | 3 | 0.99 |
| dsDNA | F3-WT | 7.38 | 3 | 0.98 |
| | F3-S2D | 5.04 | 3 | 0.99 |
| | F3-S2A | 3.64 | 3 | 0.98 |
| NCP | F3-WT | 16.69 | 7 | 0.92 |
| | F3-S2D | 14.64 | 7 | 0.92 |
| | F3-S2A | 4.56 | 7 | 0.95 |

**Figure S3.** Related to Figure 3. **S277/282 regulate RAD51AP1 binding to ssDNA, dsDNA, and the NCP.** *A*, SDS-PAGE to show purified MBP-F3-His$_6$ and mutants (1 µg protein each). *B*, Schematic representation of the RAD51AP1 protein (isoform 2; black) and the RAD51AP1-F3 fragment (residues 188-335; green) containing the bipartite DNA binding domain (grey) and the critical residues engaged in DNA bindings and previously identified in this domain(5). Location of residues S277 and S282 is indicated. *C*, Representative EMSA showing the mobility shifts for MBP-F3-His$_6$ and mutants with ssDNA. *D*, Determination of the apparent binding constants ($K_{D(app)}$) for ssDNA (n = 3-4; bars represent the means ± 1 SD). *E*, Representative EMSA showing the mobility shifts for MBP-F3-His$_6$ and mutants with dsDNA. *F*, Determination of the apparent binding constants ($K_{D(app)}$) for dsDNA (n = 3; bars represent the means ± 1 SD). *G*, Representative EMSA showing the mobility shifts for MBP-F3-His$_6$ and mutants with the NCP. *H*, Determination of the apparent binding constants ($K_{D(app)}$) for the NCP (n = 3; bars represent the means ± 1 SD). *I*, Summary of apparent $K_D$ values determined by fitting the binding data to a one-site specific binding model using non-linear regression in GraphPad Prism 10.

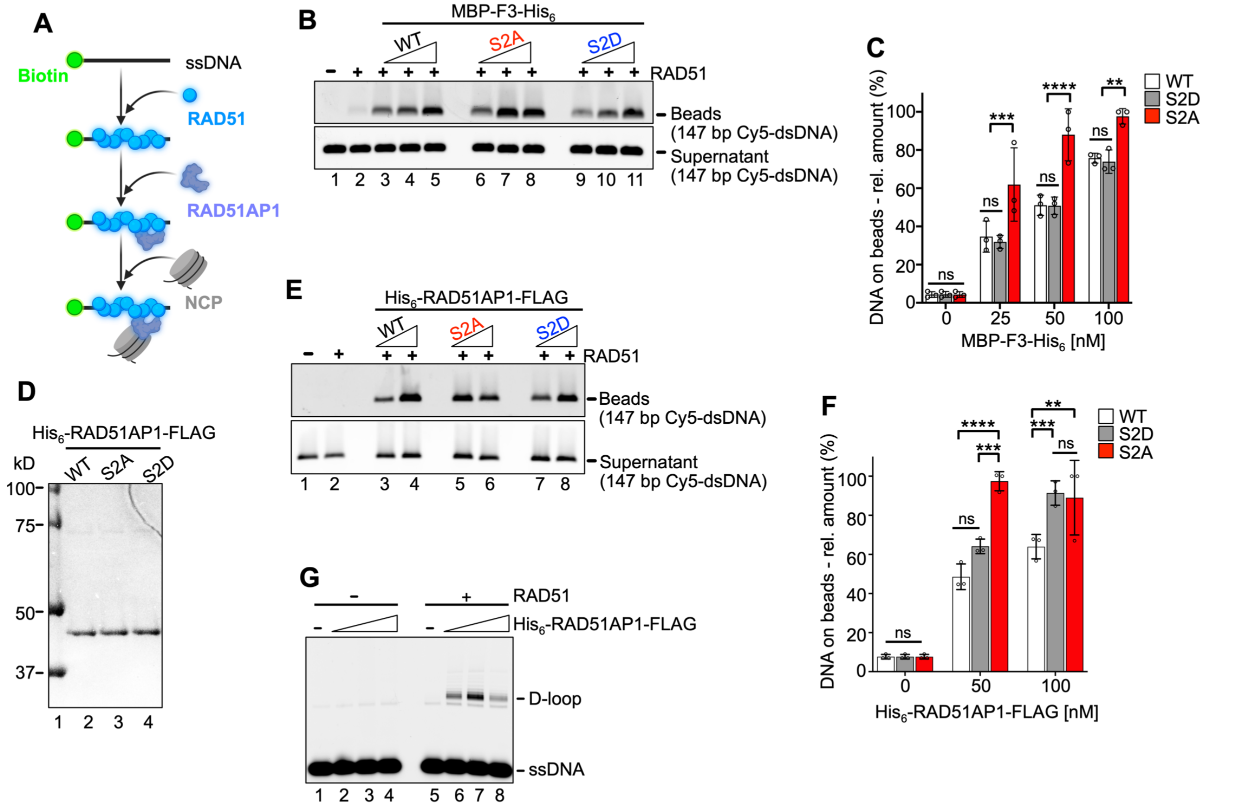


**Figure S4.** Related to Figure 4. **S277/282 regulate RAD51AP1's stimulation of RAD51 in duplex capture with the NCP.** *A*, Schematic of the duplex capture assay with His$_6$-RAD51AP1-FLAG (or mutants) and the NCP. *B*, Representative agarose gel to quantify duplex capture (*i.e.*, DNA on beads) with MBP-F3-His$_6$ or mutants and the NCP. *C*, Quantitation of duplex capture with MBP-F3-His$_6$ or mutants (n = 3; bars represent the means ± 1 SD). *D*, SDS-PAGE to show purified His$_6$-RAD51AP1-FLAG and mutants (250 ng protein each). *E*, Representative agarose gel to quantify duplex capture (*i.e.*, DNA on beads) with full-length His$_6$-RAD51AP1-FLAG or mutants and the NCP. *D*, Quantitation of duplex capture with full-length His$_6$-RAD51AP1-FLAG or mutants (n = 3; bars represent the means ± 1 SD). *G*, Agarose gel to show that the stimulation of D-loop formation by RAD51AP1 (50-200 nM) is dependent on RAD51 (lanes 6-8). Two-way ANOVA followed by Tukey's multiple comparison test. $^{**}p < 0.01$. $^{***}p < 0.001$. $^{****}p < 0.0001$. ns, not significant.

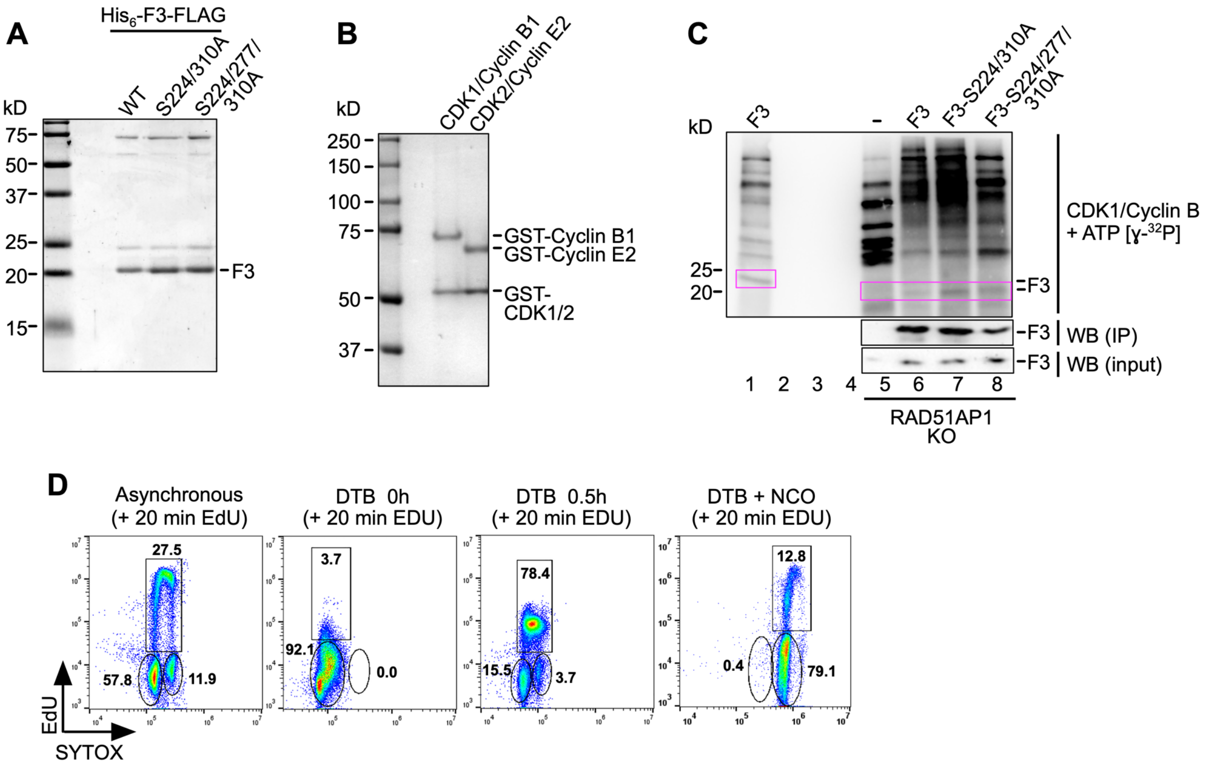


**Figure S5.** Related to Figure 5. **S277 is a CDK2 target in cells.** *A*, SDS-PAGE to show purified F3 and mutants (250 ng protein each). *B*, SDS-PAGE to show commercially obtained CDK1/Cyclin B1 (Abcam; ab271456) and CDK2/Cyclin E2 complexes (Abcam; ab268396) (500 ng protein complex each). *C*, Results from kinase reaction of transiently expressed (in HeLa RAD51AP1 KO cells) and precipitated FLAG-F3 mutants (S224/310A and S224/277/310A) with CDK1/Cyclin B1 and ɣ-$^{32}$P-ATP. Upper: Phosphorimage; middle and lower: Western blot with α-FLAG antibody. The product of the kinase reaction with purified (from *E. coli*) $His_6$-F3-FLAG is shown for comparison purposes (lane 1). *D*, Representative results from flow cytometry showing two-color fluorescence of cell cycle profiles of asynchronous RAD51AP1 KO cells stably expressing RAD51AP1, and of these cells synchronized by a DTB and released into regular growth medium without and with NCO. Prior to fixation, cells were pulse-labeled by EdU. Y-axis: EdU, X-axis: SYTOX.

**Table S1. List of oligonucleotides/primers used in this study[1].**

| Template/ plasmid | Oligonucleotide | Sequence (5´-3´) |
|---|---|---|
| FLAG-RAD51AP1 into pENTR1A (from pOK24 (1)) | 5´-FLAG forward | ATCGGTCGACCCACCATGGATTACAAAGACGATGACGATAAAGGTGTGCGGCCTGTGAGACATAAGAAACC |
| | reverse | ATCGGTACCTCAGGTGCTAGTGGCATTTGGATGCAAAG |
| FLAG-RAD51AP1-S282A (in pENTR1A) | S282A forward | TCAGCTGAAgcaAAGAAACCTAAATGG |
| | S282A reverse | AGGGTCGCGTATTTCTAATG |
| FLAG-RAD51AP1-S277/282A (in pENTR1A) | S277A forward | AGAAATACGCgcaCCTTCAGCTG |
| | S277A reverse | AATGGTTTCCTAGTGGTATC |
| FLAG-RAD51AP1-S282D (in pENTR1A) | S282D forward | GCTGAAgacAAGAAACCTAAATGGGTC |
| | S282D reverse | TGAAGGGTCGCGTATTTCTAATGGTTTC |
| FLAG-RAD51AP1-S277/282D (in pENTR1A) | S277D forward | GctgAAAGCAAGAAACCTAAATGGGTC |
| | S277D reverse | TGAAGGGATCGCGTATTTCTAATGGTTTC |
| $His_6$-RAD51AP1-S282A-FLAG (in $His_6$-RAD51AP1-FLAG /pQE-80L (6)) | S282A forward | TCAGCTGAAgcaAAGAAACCTAAATGG |
| | S282A reverse | AGGGTCGCGTATTTCTAATG |
| $His_6$-RAD51AP1-S277/282A-FLAG (in $His_6$-RAD51AP1-FLAG/ pQE-80L (6)) | S277A forward | AGAAATACGCgcaCCTTCAGCTG |
| | S277A reverse | AATGGTTTCCTAGTGGTATC |
| $His_6$-RAD51AP1-S282D-FLAG (in $His_6$-RAD51AP1-FLAG/ pQE-80L (6)) | S282D forward | GCTGAAgacAAGAAACCTAAATGGGTC |
| | S282D reverse | TGAAGGGTCGCGTATTTCTAATGGTTTC |
| $His_6$-RAD51AP1-S277/282D-FLAG (in $His_6$-RAD51AP1-FLAG/ pQE-80L (6)) | S277D forward | GctgAAAGCAAGAAACCTAAATGGGTC |
| | S277D reverse | TGAAGGGATCGCGTATTTCTAATGGTTTC |

| | | |
|---|---|---|
| MBP-F3-S282A-$His_6$ (in MBP-F3-$His_6$/ pET24a (7)) | S282A forward | TCAGCTGAAgcaAAGAAACCTAAATGG |
| | S282A reverse | AGGGTCGCGTATTTCTAATG |
| MBP-F3-S277/282A-$His_6$ (in MBP-F3-$His_6$/ pET24a (7)) | S277A forward | AGAAATACGCgcaCCTTCAGCTG |
| | S277A reverse | AATGGTTTCCTAGTGGTATC |
| MBP-F3-S282D-$His_6$ (in MBP-F3-$His_6$/ pET24a (7)) | S282D forward | GCTGAAgacAAGAAACCTAAATGGGTC |
| | S282D reverse | TGAAGGGTCGCGTATTTCTAATGGTTTC |
| MBP-F3-S277/282A-$His_6$ (in MBP-F3-$His_6$/ pET24a (7)) | S277D forward | GgctGAAGACAAGAAACCTAAATGGGTC |
| | S277D reverse | TGAAGGATCGCGTATTTCTAATGGTTTC |
| $His_6$-F3-FLAG (from $His_6$-RAD51AP1-FLAG/pQE-80L (6)) | F3 forward | ATCGGGATCCGATTCTGAGGATGATTCTG |
| | F3 reverse | CGATGTCGACTCATTTATCGTCATCGTCTTTGTAATCACCGGTGCTAGTGGCATTTGGATGCAAAGG |
| $His_6$-F3-S310A-FLAG (in pQE-80L) | S310A forward | AGAAATACGCgcaCCTTCAGCTG |
| | S310A reverse | AATGGTTTCCTAGTGGTATC |
| $His_6$-F3-S224/310A-FLAG (in pQE-80L) | S224A forward | GAAGGTAAAAgccCCAGTAGAAAAG |
| | S224A reverse | ACTTCTTTCTTTTTAATTTCTTTAACTTTAC |
| $His_6$-F3-S224/277/310A-FLAG (in pQE-80L) | S277A forward | AGAAATACGCgcaCCTTCAGCTG |
| | S277A reverse | AATGGTTTCCTAGTGGTATC |
| FLAG-F3 in pcDNA3.1 (from His6-F3-FLAG/pQE-80L) | forward | ATCGGCGGCCGCCCACCATGGATTACAAAGACGATGACGATAAAGGTGATTCTGAGGATGATTCTGATTTTTGTGAG |
| | reverse | CGATGGATCCTCAGGTGCTAGTGGCATTTGGATGCAAAG |

| | | |
|---|---|---|
| Cy5-147 bp (from pUC19-601; (8)) | 5´Cy5-147bp forward | ATATCTGAGAATCCGGTGCCG |
| | 147bp reverse | GGATGTATATATCTGACACGTGCCTG |
| biotin-ssDNA | 80-mer | TCGTAGACAGCTCTAGCACCGCTTAAACGCACGTAGGCGCTGTCCCCCGCGTTTTAACCGCCAAGGGGATTACTCCCTAG |
| Cy5-ssDNA | 80-mer | TCGTAGACAGCTCTAGCACCGCTTAAACGCACGTAGGCGCTGTCCCCCGCGTTTTAACCGCCAAGGGGATTACTCCCTAG |
| Cy5-ssDNA | 90-mer | AAATCAATCTAAAGTATATATGAGTAAACTTGGTCTGACAGTTACCAATGCTTAATCAGTGAGGCACCTATCTCAGCGATCTGTCTATTT |

[1]Mutated bases in lower case character.
